# White Matter Hyperintensities Precede other Biomarkers in *GRN* Frontotemporal Dementia

**DOI:** 10.1101/2025.03.31.646450

**Authors:** Mahdie Soltaninejad, Mahsa Dadar, D. Louis Collins, Reza Rajabli, Vikram Venkatraghavan, Arabella Bouzigues, Lucy L. Russell, Phoebe H. Foster, Eve Ferry-Bolder, John C. van Swieten, Lize C. Jiskoot, Harro Seelaar, Raquel Sanchez-Valle, Robert Laforce, Caroline Graff, Daniela Galimberti, Rik Vandenberghe, Alexandre de Mendonça, Pietro Tiraboschi, Isabel Santana, Alexander Gerhard, Johannes Levin, Benedetta Nacmias, Markus Otto, Maxime Bertoux, Thibaud Lebouvier, Chris R. Butler, Isabelle Le Ber, Elizabeth Finger, Maria Carmela Tartaglia, Mario Masellis, James B. Rowe, Matthis Synofzik, Fermin Moreno, Barbara Borroni, Jonathan D. Rohrer, Yasser Iturria Medina, Simon Ducharme, GENFI

## Abstract

**INTRODUCTION:** Increased white matter hyperintensities (WMHs) have been reported in genetic frontotemporal dementia (FTD) in small studies, but the sequence of WMH abnormalities relative to other biomarkers is unclear.

**METHODS:** Using a large dataset (n=763 GENFI2 participants), we measured WMHs and examined them across genetic FTD variants and stages. Cortical and subcortical volumes were parcellated, and serum neurofilament light chain (NfL) levels were measured. Biomarker progression was assessed with discriminative event-based and regression modeling.

**RESULTS:** Symptomatic *GRN* carriers showed elevated WMHs, primarily in the frontal lobe, while no significant increase was observed in *C9orf72* or *MAPT* carriers. WMH abnormalities preceded NfL elevation, ventricular enlargement, and cortical atrophy. Longitudinally, baseline WMHs predicted subcortical changes, while subcortical volumes did not predict WMH changes, suggesting WMHs may precede neurodegeneration.

**DISCUSSION:** WMHs are elevated in a subset of *GRN*-related FTD. When present, they appear early and should be considered in disease progression models.

**Highlights:**

- Elevated WMH volumes in symptomatic *GRN* carriers, but not in other mutations.
- WMH accumulation is mostly observed in the frontal lobe.
- WMH abnormalities appear early in *GRN*-FTD, before NfL, atrophy, and ventriculomegaly.
- Longitudinally, WMH volumes can predict subcortical changes, but not vice versa.
- WMHs are key early markers in *GRN*-FTD and should be included in progression models.

**RESEARCH IN CONTEXT:** *Systematic review:* We systematically reviewed the literature on white matter hyperintensities (WMHs) in frontotemporal dementia (FTD) using PubMed. While a few small studies reported increased WMHs in *GRN* mutation carriers, their sample sizes were limited, and they did not assess the timing of WMHs within disease progression or their temporal relationship to other biomarkers.

***Interpretation**:* We identified a sequence of key biomarkers in *GRN*-related FTD and demonstrated that WMHs are among the earliest biomarkers, preceding cortical and subcortical atrophy as well as blood biomarkers. This aligns with neuropathological evidence of early white matter involvement in FTLD-*GRN*. Additionally, using a larger dataset, we validated previous reports of elevated WMHs in *GRN* carriers, confirming their reliability.

***Future directions**:* Future studies should integrate WMHs into FTD progression models to enhance early diagnosis. Understanding why only a subset of *GRN* carriers exhibit high WMH volumes remains a key research priority.

## 1 BACKGROUND

Frontotemporal dementia (FTD) presents as a multifaceted neurodegenerative disorder, marked by progressive deterioration in behavior, personality, and/or language, ranking as the second most prevalent cause of early onset dementia following Alzheimer’s disease.^1^ Approximately 30% of FTD cases exhibit a robust familial dementia history, often linked to specific genetic mutations. The majority of FTD’s heritability stems from autosomal dominant mutations within three primary genes: *C9orf72* (chromosome 9 open reading frame 72), *GRN* (progranulin), and *MAPT* (microtubule-associated protein tau).^2^ Despite symptomatic overlap across these gene mutations, the molecular mechanisms driving the emergence of phenotypic outcomes are inherently distinct.

White matter hyperintensities (WMHs) have garnered significant attention due to their clinicopathologic contributions in a range of neurodegenerative conditions and their deleterious effect on cognition.^3,4^ These lesions manifest as hyperintensities on specific sequences of magnetic resonance imaging (MRI), signaling abnormalities within the brain’s white matter. These anomalies can indicate areas of demyelination, gliosis, and/or small vessel disease. WMH post mortem histopathology reveals non-specific brain alterations, such as gliosis, myelin and axon loss attributed to arteriosclerosis, tissue rarefaction, and lipohyalinosis. These changes may be caused by various factors, including hypoxia, hypoperfusion, blood-brain barrier leakage, inflammation, degeneration, and amyloid angiopathy.^5^ Moreover, increasing evidence suggests that WMHs in neurodegeneration are not solely driven by vascular pathology but may also reflect intrinsic disease processes, including amyloidosis and gray matter degeneration, as shown in Alzheimer’s disease, where WMHs have been linked to cerebral amyloid angiopathy and neurodegeneration rather than traditional vascular risk factors.^6^

The overwhelming majority of neuroimaging research in FTD has centered on changes in gray matter, with less attention given to the role of WMHs. Sudre et al.^7^ reported increased WMH burden in FTD patients with symptomatic *GRN* mutations, but not in those carrying *MAPT* or *C9orf72* mutations. This investigation did not find significant WMH changes during the presymptomatic phase. A follow-up longitudinal study from the same group showed variations among *GRN* cases with 25% of individuals displaying either no WMH or only mild WMH during the symptomatic phase, and only 9% of those in the presymptomatic phase already exhibiting severe WMH involvement.^8^ Despite its reliance on a small sample size, this study has exerted a significant influence on the research landscape concerning the association between FTD and WMH being tied to *GRN* cases. Another study^9^ employing Diffusion Tensor Imaging revealed microstructural white matter changes among individuals with *C9orf72* repeat expansions and *MAPT* mutations during presymptomatic stages, highlighting the impact that genetic FTD has on white matter.

Despite these results, WMHs were not factored in recent disease models aiming to guide clinical trials.^10^ In our study, we aim to address these gaps in knowledge by investigating the prevalence of WMH across distinct genetic groups and different stages of FTD, leveraging a later version of the GENFI2 dataset with a larger sample to assess the validity of previous WMH studies. We further undertook a temporal analysis of WMH compared to other markers in the course of the progression of the disease.

## 2 METHODS

### 2.1 Data

Data for this study were obtained from the fifth data freeze of GENFI2 (Genetic Frontotemporal Dementia Initiative), a large international study involving 25 centers in Europe and Canada. GENFI collects longitudinal data on genetic FTD and aims to gather multimodal neuroimaging, cognitive, and fluid biomarkers to develop markers for early-stage FTD identification, track disease progression, and gain insight into the presymptomatic phase of the disease. This study received the approval of the McGill University Health Centre review ethics board (MP-20-2016-2500), and all data-collecting sites obtained their local board approval. Written informed consent was obtained from all participants, and the research was conducted according to the ethical principles outlined in the Declaration of Helsinki.

Participants included in GENFI2 were known symptomatic carriers of pathogenic mutations in *C9orf72*, *GRN*, or *MAPT*, as well as their first-degree relatives who were at risk of carrying a mutation. Genotyping was performed at local sites, and all participants underwent a standard clinical evaluation including medical and family history assessments, as well as physical examinations. Symptomatic carriers met clinical criteria for behavioral variant of Frontotemporal Dementia (bvFTD), Primary Progressive Aphasia (PPA), Frontotemporal Dementia with Amyotrophic Lateral Sclerosis (FTD-ALS) or other rare presentations, while presymptomatic carriers did not fulfill clinical criteria. The non-carrier group comprised healthy first-degree relatives of symptomatic carriers who tested negative for the reported family mutation. Detailed inclusion and exclusion criteria can be found elsewhere.^11^

MRI scans were acquired using 3T scanners. Serum levels of NfL (neurofilament light chain) and GFAP (glial fibrillary acidic protein) were longitudinally measured using the single-molecule array technique (Simoa).^12^

GENFI is a longitudinal dataset with multiple visits available for some participants. In this study, we first used a cross-sectional design to maximize the number of subjects, selecting only one visit per participant (except in our final analysis, which incorporated a longitudinal approach). To ensure consistency in data selection, we prioritized visits with both blood and imaging biomarkers available. Where participants had multiple eligible visits, we chose the most recent one to better capture a broader spectrum of disease severity. Longitudinal visits were used to test for the temporal relationship of biomarker changes.

### 2.2 Image Processing

#### 2.2.1 White Matter Hyperintensities Segmentation

The segmentation of WMH was performed using BISON,^13^ integrating data from T1-weighted (T1w) and T2-weighted (T2w) imaging modalities. The workflow employed a random forest classifier trained with location, intensity parameters, and manually labeled data to generate participant-specific WMH maps. Before WMH segmentation, T1w and T2w scans underwent preprocessing steps including image denoising, intensity non-uniformity correction, and intensity normalization within a range of 0-100. T1w images were linearly registered and subsequently nonlinearly registered to the ICBM152 template. T1w and T2w images were linearly co-registered using a six-parameter rigid registration. All stereotaxic space priors and averages for WMH segmentation were resampled onto the native T1w volume using the inverse of the estimated non-linear registration transformation. Quality control was performed by visually evaluating WMH segmentations using Qrater.^14^ As a result, 11 out of 793 cases, including outliers and poorly segmented data, were excluded (Fig. 1). The WMH lesion maps were then linearly registered to the ICBM152 template, and Hammers’ atlas was used to quantify WMH volumes in each of the eight lobes.^15,16^

**Figure 1.**
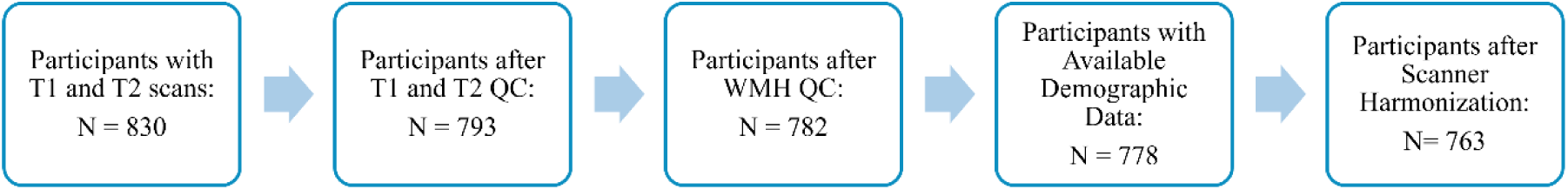
Flowchart of inclusion and exclusion criteria in the study. QC: Quality control; WMH: White matter hyperintensity.

#### 2.2.2 Brain Parcellation

In this study, gray matter volumes were obtained with the Geodesic Information Flow (GIF) algorithm,^17^ a multi-atlas segmentation approach, for accurate and robust cortical and subcortical volume parcellation. GIF utilizes spatially-variant graph structures, connecting morphologically similar participants for gradual information diffusion amid large-scale morphological variability. From the parcellated regions, we considered the volumes of cortical and subcortical areas most closely associated with FTD, as identified in previous studies,^18–22^ including the frontal lobe, temporal lobe, insula, basal ganglia (nucleus accumbens, caudate, putamen, and globus pallidus), cerebellum, cingulate cortex, ventricle, amygdala, hippocampus, and thalamus. The summed volumes of left and right regions were then utilized for further analysis. Additionally, to address individual variations in brain size, the volumes were standardized by dividing them by the intracranial volume.

### 2.3 Statistical Tests

Statistical analyses were conducted using RStudio version 4.3.1. To achieve a normal distribution of the WMH volumes, log transformation was applied. A mixed-effects model was employed to adjust the WMH volumes of mutation carriers, with age, sex, and scanner site being taken into account. The model was specified as follows:

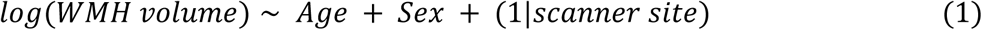

The model was fitted using data from healthy controls at each scanning site. Subsequently, the difference between the actual log-transformed WMH volumes and the predicted volumes derived from this model for each person was calculated. These differences, referred to as adjusted WMH values, capture residual WMH volume after considering demographic and technical variables.

During this stage of the analysis, 15 participants were excluded due to the unavailability of control data from their scanning site, which made data harmonization impossible (Fig. 1). Detailed information regarding the exact number of participants and their demographic characteristics is provided in Table 1.

**Table 1.**
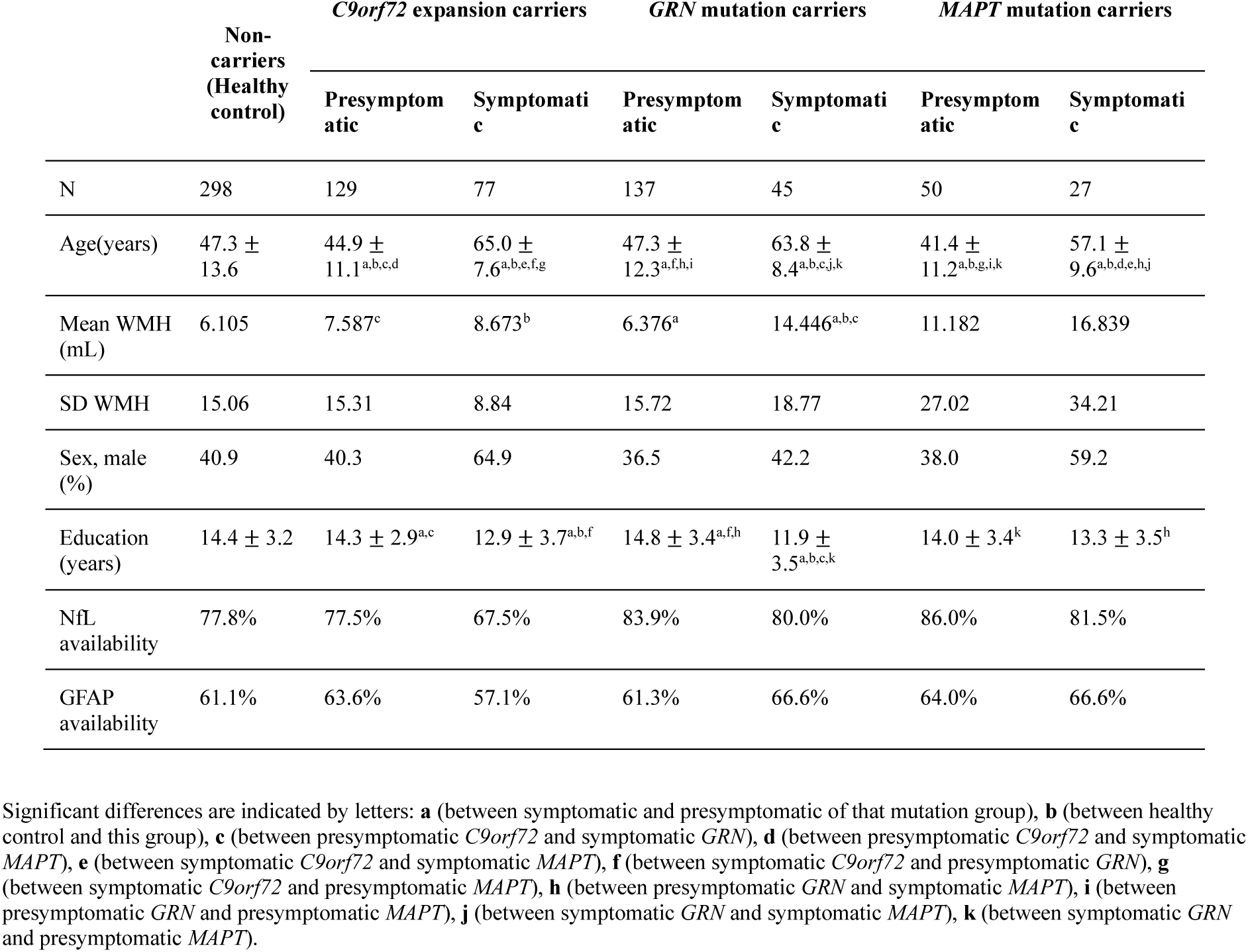
Demographic characteristics of participants.

Kruskal-Wallis variance analysis was performed to compare adjusted WMH values across the three mutation cohorts and different disease stages (symptomatic and presymptomatic). Dunn’s post-hoc test, a non-parametric pairwise multiple comparison test, was conducted to determine significant differences in adjusted WMH volume between mutation groups. A similar analysis was performed for adjusted WMH volumes in each of the eight lobes. All p-values reported in the manuscript are corrected for multiple comparisons with Bonferroni’s method.

To rule out confounding influences on WMH, cardiovascular risk factors (stroke, hypertension, hypercholesterolemia, and diabetes) and history of traumatic brain injury were compared across mutation groups and controls. Since no significant differences were found, these risk factor variables were not included as covariates in the model.

### 2.4 Temporal Relationship Analysis

After investigating the prevalence of WMH across distinct genetic groups, we aimed to determine the temporal sequence of WMH accumulation in relation to other key imaging biomarkers in groups exhibiting substantial WMH burden. In addition to lobar WMH and WMH of the whole brain, we included essential gray matter volumes and subcortical measures in FTD, i.e., frontal and temporal gray matter, cingulate, insula, cerebellum, basal ganglia (nucleus accumbens, caudate, putamen, and globus pallidus), hippocampus, amygdala, and thalamus, alongside additional biomarkers like ventricle volume, GFAP, and NFL for ranking.

#### 2.4.1 Cross-Sectional Analysis Using Discriminative Event-Based Modeling

We utilized the Discriminative Event-Based Modeling (DEBM) approach,^23,24^ to investigate the order of biomarker abnormalities in presymptomatic and symptomatic FTD. DEBM is well-suited for our purpose because it requires only cross-sectional data and effectively handles missing values. In DEBM, an “event” refers to the transition of a biomarker from a normal to an abnormal state, with the total number of events in disease progression corresponding to the number of biomarkers. To ensure a more normalized distribution, blood biomarkers (NfL and GFAP), ventricle volume and WMHs were log-transformed, effectively reducing skewness in their distributions. To control for confounding factors, the DEBM analysis incorporated sex and age by adjusting biomarker values based on these factors prior to Gaussian Mixture Modeling (GMM).

The DEBM procedure initially determines the distribution of normal and abnormal biomarker values through GMM. Using these distributions, it computes the probability for each participant that the biomarker is abnormal. This probability signifies the progression of that biomarker. Therefore, based on these probabilities, we create an approximate sequence of biomarker abnormality for each participant, which is aggregated across participants to create a robust central biomarker ordering that minimizes the sum of distances to all participant-wise orderings.

Healthy controls, presymptomatic carriers, and symptomatic carriers were treated as distinct diagnoses to effectively model the progression. The degree of uncertainty in biomarker ordering was assessed by estimating it for 100 independently sampled datasets using bootstrap resampling with replacement. The analysis included 778 participants with verified imaging biomarkers.

For the DEBM approach to be effectively applied, two key assumptions must be considered when selecting biomarkers:

1. The biomarker must exhibit a statistically significant difference between the FTD and healthy control groups.
2. The Gaussian Mixture Model must be fitted appropriately, which we verified by calculating the mean squared error of the fitted distribution for each biomarker.

We rigorously assessed the accuracy of the GMM by calculating the mean squared error and visually inspecting the distribution. The limited sample size constrained the applicability of the DEBM approach to certain biomarkers. Biomarkers that did not meet these criteria were excluded from the DEBM analysis and subsequently evaluated using the longitudinal approach. This ensured the incorporation of only reliable biomarkers into the DEBM, while allowing further investigation of excluded biomarkers through complementary methods. This integrated approach enabled us to maximize the utility of both cross-sectional and longitudinal data, providing a comprehensive understanding of biomarker dynamics.

To validate the accuracy of the model, we assessed the disease stage that the DEBM estimated for each individual. This was done by comparing the abnormality probabilities of biomarkers for each participant with the central biomarker ordering derived from the model. Two validation approaches were employed. First, we evaluated the model’s performance by calculating the area under the curve (AUC) to differentiate between symptomatic carriers and healthy controls based on estimated disease stage using 10-fold cross-validation. As a second, construct, validation metric, we examined the correlation between the estimated disease stages and key clinical scores commonly used in FTD assessment. These included the Mini-Mental State Examination (MMSE), Trail Making Test Part B (TMT-B), Boston Naming Test, Digit Symbol substitution test, and Verbal Fluency test. Nonparametric Spearman’s rank correlation analyses were conducted to assess the relationships between the estimated disease stages and key clinical scores, given the non-normal distribution of the overall population on these metrics. These validation metrics allowed us to assess both the discriminative power and clinical relevance of the estimated disease stages.

#### 2.4.2 Longitudinal Analysis Using Linear Regression Modeling

Subsequently, we conducted a longitudinal assessment focusing on the interplay between WMH and other neuroimaging biomarkers that were not evaluated through DEBM analysis, specifically in *GRN* mutation carriers. These biomarkers include the insula, basal ganglia (nucleus accumbens, caudate, putamen, and globus pallidus), thalamus, hippocampus, amygdala, and cingulate volumes. The process involved quantifying the z-scores for each biomarker and employing linear regression models to predict the shift in each biomarker between baseline and follow-up measurements, with WMH serving as either the predictor or response biomarker in each model. The models were designed to account for potential influences from age, sex, NfL, and education factors. The analysis was represented by equation 2.

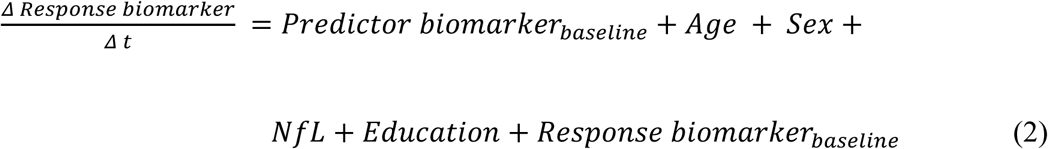

Here, 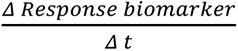 represents the predicted rate of change in the response biomarker between the baseline measurement and all subsequent follow-up evaluations. In our analysis, WMH volumes, and NfL biomarkers were log-transformed to achieve a normal distribution.

## 3 RESULTS

### 3.1 Participant Demographics and Clinical Information

Table 1 provides an overview of demographic and clinical data. The research included 298 family controls who did not carry the mutation. Among the 465 participants identified with mutations, *C9orf72* mutations were the most prevalent, affecting 44.3% of the group, followed by *GRN* mutations at 39.1% and *MAPT* mutations at 16.6%. Notably, 68% of these individuals were in the asymptomatic stage across the genetic variants mentioned. Among symptomatic participants, the predominant diagnosis was bvFTD, representing 66.4% of cases, with PPA at 16.8%, and Amyotrophic Lateral Sclerosis (ALS) or FTD-ALS at 10.7%. The remaining 6.1% of cases were diagnosed with other clinical syndromes.

Demographic analyses confirmed that symptomatic mutation carriers were older and had received fewer years of education than both presymptomatic mutation carriers and control groups (p<0.001). *MAPT* mutation carriers and the control group were younger than those with the *GRN* mutation (*MAPT*: p=0.01; controls: p=0.001) and those carrying the *C9orf72* mutation (*MAPT*: p=0.002; controls: p<0.001). Additionally, symptomatic *C9orf72* mutation carriers had fewer years of education compared to the control group (p=0.03). The gender ratio among symptomatic carriers also showed a higher proportion of males compared to the presymptomatic (p<0.001) and control groups (p=0.002). Moreover, the *C9orf72* symptomatic carrier cohort included a significantly higher proportion of males compared to *GRN* symptomatic carriers (p=0.02). Other demographic characteristics remained consistent across all groups. NfL samples were obtained from 78.64% of participants, and GFAP data were available for 61.86%.

### 3.2 White Matter Hyperintensities Across Genetic Groups

The WMH distribution maps in Fig. 2 depict the absolute prevalence and regional distribution of WMH across mutation groups on a voxel-wise basis. As expected, in all participants (including controls) WMHs are predominantly located in the periventricular regions, with a visually wider spatial extent in symptomatic participants.

**Figure 2.**
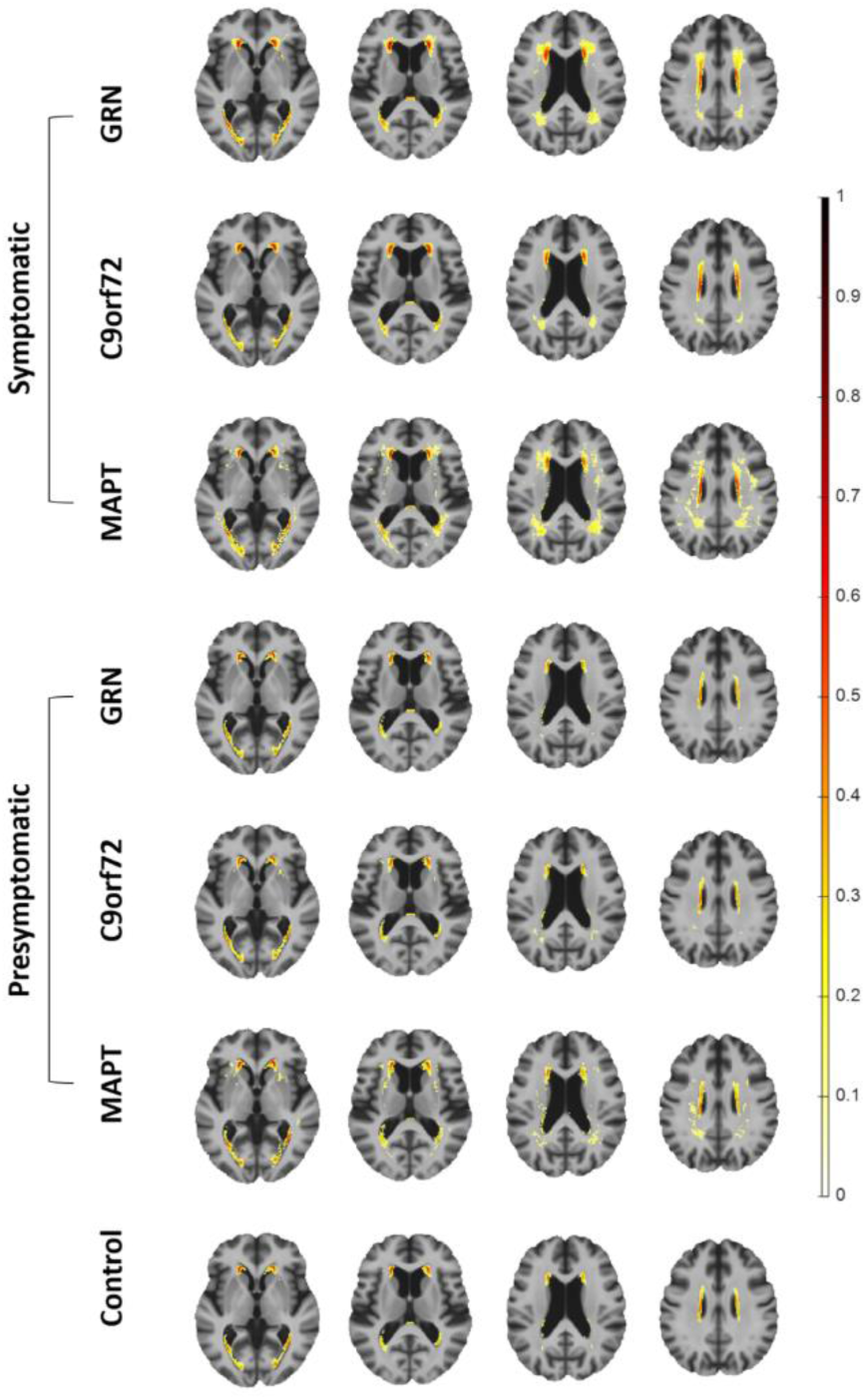
Voxel-wise distribution of WMH prevalence in mutation groups. The colour bar represents the proportion of participants within each cohort exhibiting WMHs at specific voxel locations.

Fig. 3 presents the statistical comparisons of adjusted volumes of WMH, as described in the method section. These values are log-transformed initially and then controlled for age, sex, and scanner site. For precise measures, refer to Table S1 and S2 in the supplementary materials, these values depict the contrast of each cohort as compared to the control group. Adjusted WMH volumes were significantly higher among all-group symptomatic mutation carriers as a group compared to controls (P_Bonferroni_ = 0.012). Specifically, post-hoc test showed that the effect was driven by symptomatic carriers with *GRN* mutations who exhibited markedly elevated whole brain WMH volumes compared to controls (P_Bonferroni_ < 0.001), while the difference was not present for *C9orf72* and *MAPT*. Among symptomatic cases, *GRN* carriers also showed higher WMH volumes compared to those with *C9orf72* (P= 0.016) and *MAPT* (P=0.024) mutations; however, these differences did not reach significance after Bonferroni correction. Within the *GRN* mutation carriers, a distinctive pattern emerged, as symptomatic cases exhibited higher WMH volumes than presymptomatic *GRN* cases (P_Bonferroni_ = 0.001), underscoring the progressive nature of WMH accumulation over the disease course. Aside from the *GRN* mutation carriers, no other significant differences in adjusted WMH volumes were observed between other mutation groups and controls or presymptomatic carriers.

**Figure 3.**
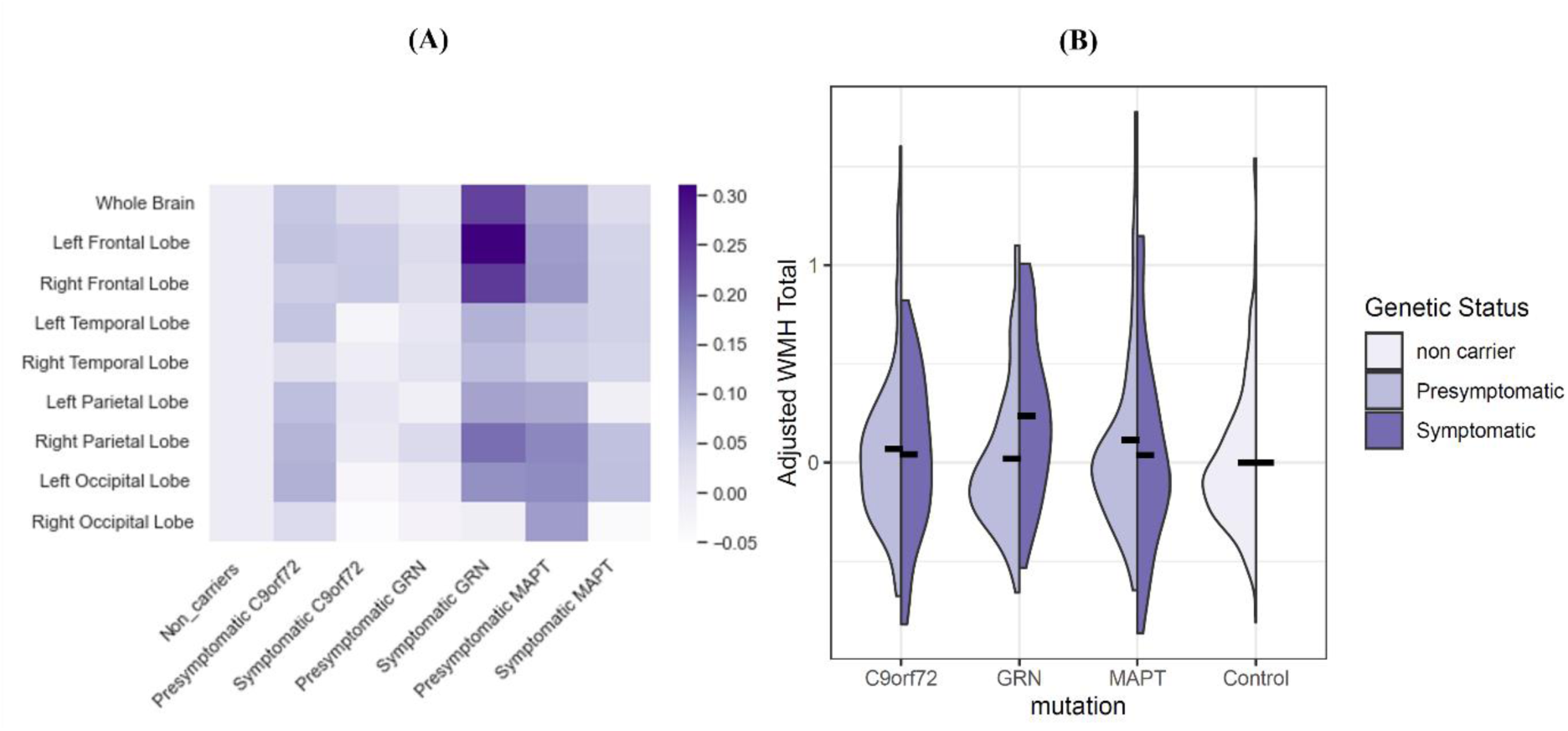
Comparison of adjusted WMH volumes across mutation cohorts. **(A)** Regional WMH volume differences across mutation cohorts, controlling for age, sex and scanner site. The values are log transformed. **(B)** Comparison of adjusted WMH volumes among different mutation cohorts. The black lines represent the average WMH volume for each group.

Examination of WMH volumes per lobe revealed that the frontal lobe was the most prominent site of accumulation. Notably, symptomatic carriers demonstrated a marked increase in adjusted WMHs compared to healthy controls in both the left frontal lobe (P_Bonferroni_ < 0.001) and the right frontal lobe (P_Bonferroni_ = 0.007). Symptomatic *GRN* carriers exhibited substantially elevated WMHs in the left frontal lobe compared to controls (P_Bonferroni_ < 0.001), symptomatic *C9orf72* (P_Bonferroni_ = 0.010), and symptomatic *MAPT* carriers (P_Bonferroni_ = 0.025). Additionally, WMH volumes in the left frontal lobe were significantly higher in symptomatic *GRN* carriers than in their presymptomatic counterparts (P_Bonferroni_ < 0.001). The pattern was also present in the right frontal lobe, where symptomatic *GRN* mutation carriers exhibited heightened WMHs compared to controls (P_Bonferroni_ = 0.003) and presymptomatic *GRN* carriers (P_Bonferroni_ = 0.009). There was an unexpected trend for high WMHs in the left occipital lobe of presymptomatic individuals across all genetic groups compared to controls; however, after adjusting p-values, this effect did not reach significance (P_Bonferroni_ = 0.171). A similar non statistically significant trend was observed for the right occipital lobe in presymptomatic carriers compared to symptomatic carriers (P_Bonferroni_ = 0.097), especially in *C9orf72* mutation carriers (P_Bonferroni_ = 0.125).

The number of cases per subtype of GRN mutation was too small to compare prevalence across them, but we provide the adjusted level of WMH per mutation subtype in supplementary figure S1.

### 3.3 Biomarker Dynamics in *GRN* Cohort

#### 3.3.1 Temporal Cascade of biomarker abnormalities

Since *GRN* mutation carriers were the only group with a significant amount of white matter hyperintensities, particularly in the frontal lobe, we focused our analysis on this cohort. We examined the temporal relationships among WMHs in the frontal lobe, WMHs in the temporal lobe, total WMHs, and other key neuroimaging biomarkers in FTD. These included frontal and temporal gray matter, insula, cerebellum, basal ganglia, thalamus, hippocampus, amygdala, and cingulate, as well as additional biomarkers including ventricle volume, GFAP, and NfL. All of these biomarkers exhibited significant differences between FTD and control groups (as reported in Table S6). Of these, nine biomarkers met the criteria for DEBM analysis, which required both significant group differences and adequate Gaussian Mixture Model (GMM) fitting as assessed by the mean squared error of the GMM distributions (Table S5). These nine biomarkers included WMHs in the frontal lobe, WMHs in the temporal lobe, total WMHs, ventricle volume, cerebellum, frontal and temporal gray matter, and levels of GFAP and NfL. The DEBM analysis was performed on this subset of biomarkers to delineate their sequence of abnormalities and associated uncertainties in *GRN* FTD, as depicted in Fig. 4. This variability was measured through 100 bootstrapping iterations (with replacement).

**Figure 4.**
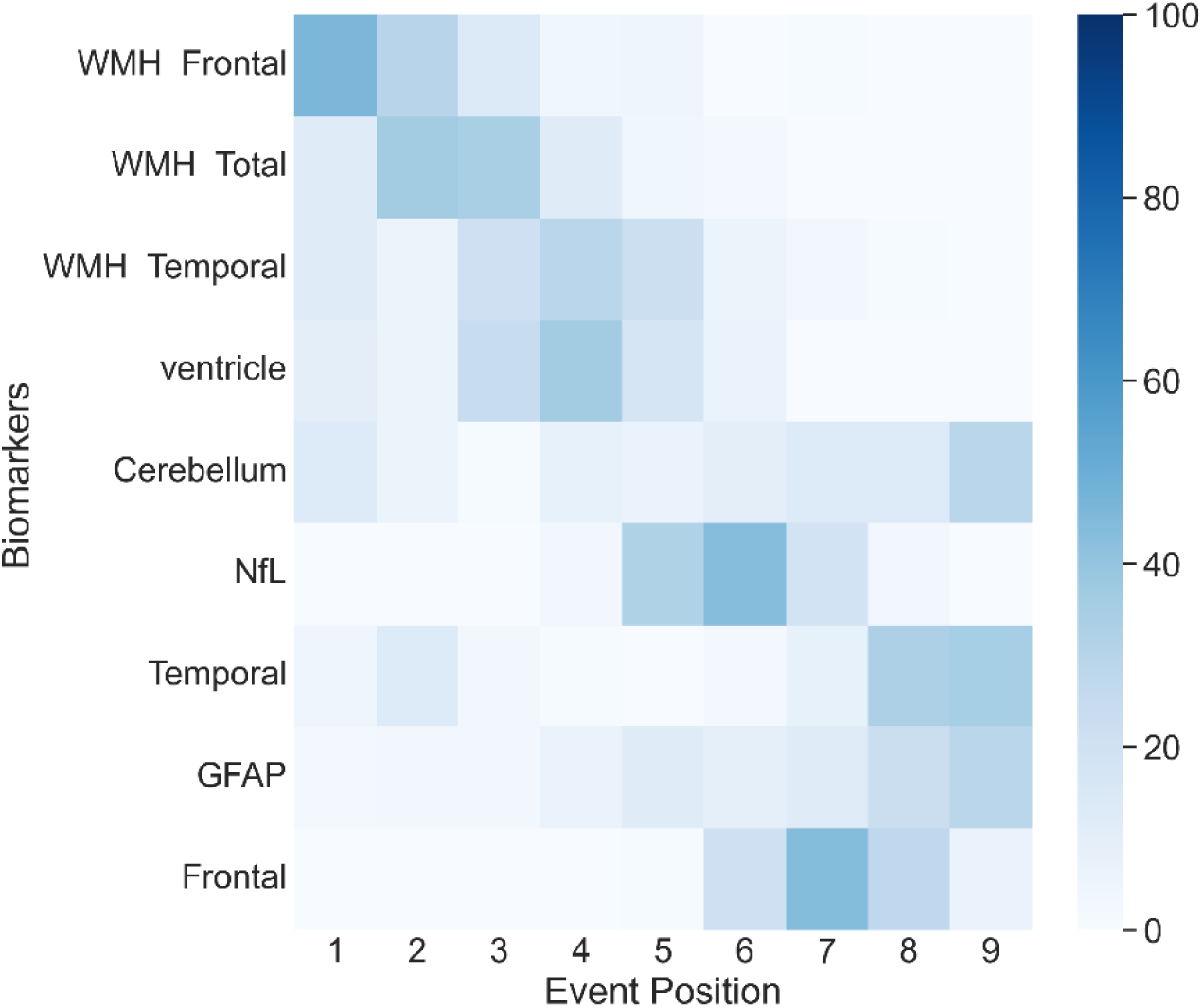
Sequence of biomarker abnormalities. The positional variance diagram for the *GRN* cohort, illustrates the most probable sequence of biomarker abnormalities along with their corresponding uncertainties. The y-axis (from top to bottom) orders the biomarkers by the most likely sequence as estimated by the DEBM model, while the x-axis indicates the position of each biomarker in the sequence, ranging from one to the total number of biomarkers. The colour intensity of each square represents the frequency with which a biomarker was placed at a specific position during bootstrap resampling. The spread from bootstrap resampling reflects the standard error of the distribution, representing the uncertainty in the estimated ordering. WMH = white matter hyperintensities; NfL = neurofilament light chain; GFAP = glial fibrillary acidic protein.

According to the findings depicted in Fig. 4, WMH abnormalities emerged at earlier disease stages compared to other studied neuroimaging biomarkers. This initial phase of WMH changes was followed by abnormalities in ventricular size and NfL levels, which were subsequently succeeded by gray matter atrophy in the temporal and frontal lobes.

#### 3.3.2 Validation of Disease Stage Estimation

The accuracy of estimated event ordering, validated through disease stage differentiation between symptomatic carriers and healthy controls, demonstrates robust clustering ability. This was reflected in high area under the curve (AUC) values for distinguishing controls from *GRN* mutation carriers, with an AUC of 0.92 ± 0.05. Further validation is provided by the strong correlation between estimated disease stages and clinical scores (including MMSE, TMT-B, Boston Naming Test, Digit Symbol, and VF), as detailed in Table S7.

### 3.4 Longitudinal Study in *GRN* Mutation Carriers

To further investigate the temporal relationships among biomarkers that were not included in the DEBM analysis, we conducted a longitudinal assessment focusing on *GRN* mutation carriers. This analysis explored the dynamic interplay between WMHs and other neuroimaging biomarkers, including the insula, basal ganglia (nucleus accumbens, caudate, putamen, and globus pallidus), thalamus, hippocampus, amygdala, and cingulate volumes. By quantifying z-scores for each biomarker and employing linear regression models, we assessed whether changes in WMH predicted alterations in these subcortical regions or vice versa over time. The longitudinal analysis encompassed 98 participants (82 presymptomatic and 16 symptomatic *GRN* carriers) who had follow-up scans, enabling us to better understand how WMH changes correlate with downstream neurodegeneration.

Estimated parameters of each model are reported in Table S8 and S9. Fig. 5A displays the T statistics for all associations, regardless of significance, among the neuroimaging biomarkers identified in our analysis. Fig. 5B focuses on significant associations (FDR corrected p-value <0.05), where a line connecting the baseline of a predictor biomarker on the left to the rate of change of a response biomarker on the right indicates that the predictor biomarker has predictive value for explaining variations in the response biomarker over time.

**Figure 5.**
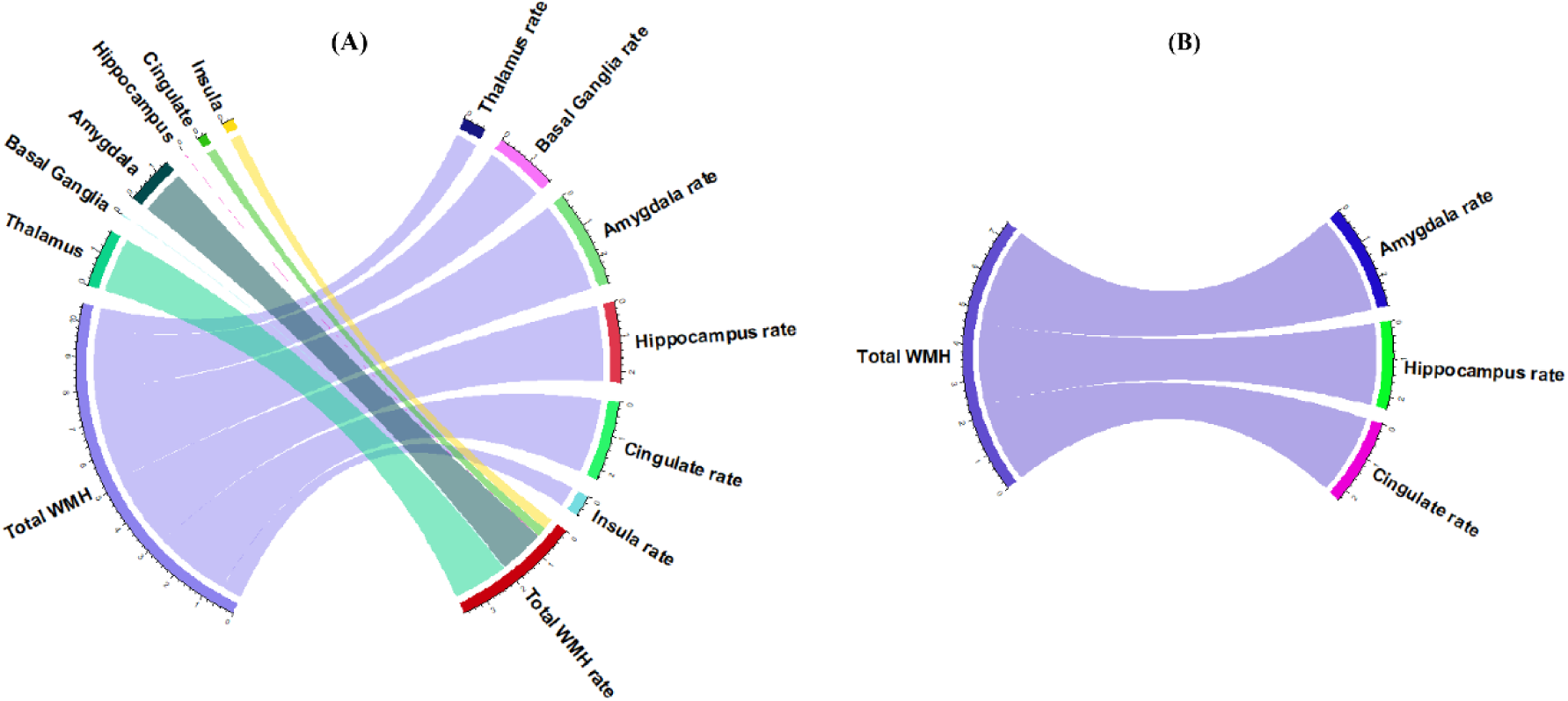
Associations between WMH and subcortical biomarkers and predictability of longitudinal variations. Diagrams illustrate the associations between predictor biomarkers (left side) and the rate of change of response biomarkers (right side). The width of the connecting lines represents the t-statistic, indicating the strength of the predictive association. **(A)** Chord diagram showing the t-statistics for all tests, including both significant and non-significant associations. **(B)** Diagram displaying only the significant associations (FDR-corrected p-value < 0.05). The figure is generated by ‘circlize’ package of R.^25^

The results shown in Fig. 5B indicate that baseline WMH volumes predict subsequent brain changes. Higher baseline WMH volumes were associated with a more rapid decrease in amygdala volume (p = 0.006, FDR-corrected p = 0.038), accelerated hippocampal atrophy (p = 0.023, FDR-corrected p = 0.050), and faster cingulate volume reduction (p = 0.025, FDR-corrected p = 0.050). Interestingly, none of the studied subcortical biomarkers (thalamus, basal ganglia, amygdala, hippocampus, cingulate and insula) were found to predict variations in WMH volumes. Our results underscore that WMHs are significant predictors of greater gray matter loss over time, emphasizing the impactful role of WMHs on brain structure changes.

## 4 DISCUSSION

Our investigation into WMH across genetic groups in the context of FTD reveals intriguing patterns. Notably, WMH prevalence is elevated in symptomatic genetic FTD due to *GRN* mutation, but not in the other mutation groups when accounting for age and sex. The spatial distribution of WMH is concentrated in the frontal lobes. Interestingly, while not present in all patients, the occurrence of WMH follows a consistent temporal pattern, emerging early in the disease progression sequence and preceding temporal and frontal cortical volume changes.

Our results underscore the specific association of WMH with the *GRN* mutation in genetic FTD. This finding is in line with Sudre et al.,^7,8^ which also observed a similar association in a study conducted on the previous smaller GENFI data release. Patients with *GRN* mutations are deficient in progranulin, a protein that plays a crucial role in regulating the growth and survival of brain cells. There is a well documented link between progranulin deficiency and neuroinflammation,^26,27^ and neuroinflammation is implicated in the pathogenesis of WMH.^28^ This observation prompts a deeper exploration of how neuroinflammation contributes to the formation of WMH in the context of *GRN* FTD, providing a potential avenue for targeted therapeutic interventions.

Our DEBM analysis elucidates a sequential pattern of biomarker abnormalities in *GRN* related FTD, beginning with abnormalities in WMHs. These early indicators of disease progression are subsequently followed by ventricular abnormalities and alterations in NfL levels, eventually leading to measurable temporal and frontal gray matter atrophy in later stages. This trajectory highlights the pivotal role of WMHs in understanding *GRN* FTD progression.

Our longitudinal analysis of *GRN* carriers reveals a noteworthy association between the initial volume of WMHs and subsequent reductions in gray matter volume across several critical brain regions, including the amygdala, hippocampus, and cingulate. This observation underscores the association between baseline WMHs and future neurological deterioration, characterized by pronounced atrophy within these areas. Conversely, our study found no evidence to suggest that alterations in subcortical biomarkers could serve as predictors for changes in WMH volume. Given the directional relationship observed, our study suggests that WMHs might precede the atrophy of gray matter, a conclusion that finds resonance in what was found in several studies of Alzheimer’s disease.^29–31^

Furthermore, our analysis indicates that NfL abnormalities precede frontal and temporal lobe atrophy, echoing the findings of Panman et al.^24^ and Staffaroni et al.^10^, who identified NfL as an early abnormal biomarker in the *GRN* mutation group, preceding changes in gray matter volumes, white matter microstructures, and cognitive markers. While Panman et al.^24^ identified NfL as the earliest biomarker among the key markers they investigated in FTD, their study did not include WMH. In contrast, our findings suggest that WMHs may precede even NfL abnormalities, indicating that WMHs could represent the earliest detectable biomarker in *GRN*-related FTD.

We also observed that abnormalities in GFAP manifest in the late stages of the disease, while abnormalities in NfL appear earlier. This finding is consistent with the sequence of fluid biomarkers in FTD reported by Van Der Ende et al.^32^

The findings of Planche and colleagues,^22^ who used lifespan brain atrophy models in FTD based on age stages, showed that subcortical atrophy precedes focal cortical atrophy within specific behavioral and/or language networks. Their work highlighted the amygdala as one of the earliest subcortical regions affected in behavioral variant FTD and semantic dementia. Our longitudinal analysis, however, suggests that WMHs can become abnormal before changes occur in subcortical regions such as the amygdala. Considering the work of Planche et al.,^22^ this implies that WMHs may serve as the earliest biomarker in behavioral variant FTD and semantic dementia.

The current paradigm for modeling disease progression in genetic FTD due to *GRN* mutation may benefit from integrating WMH into existing frameworks. Our DEBM and longitudinal analysis findings indicate early white matter disruption in FTD. A prior study by McKenna et al.^33^ highlighted alterations in white matter as relatively precise and early radiological markers, particularly effective in differentiating pre-symptomatic mutation carriers from healthy controls. Incorporating WMH into disease models may offer a more comprehensive understanding of the progression from presymptomatic to symptomatic stages, shedding light on the nuanced temporal dynamics of WMH accumulation in the context of genetic FTD.

Our findings align with neuropathological evidence showing early white matter involvement in FTLD-*GRN*. A recent study found severe frontal myelin loss in *GRN* mutation carriers, independent of axonal degeneration, suggesting a primary myelin defect.^34^ This supports our observation that WMHs appear earlier than cortical atrophy and NfL changes. Their findings also implicate microglial dysfunction and TMEM106B pathology, highlighting distinct pathogenic mechanisms and reinforcing the value of white matter biomarkers in disease staging and therapy monitoring.

It is essential to acknowledge certain limitations in our study. One limitation is that we used cortical volume as a measure of gray matter atrophy, which might not be as sensitive to subtle gray matter changes as cortical surface-based measures. Additionally, the age difference between groups, where presymptomatic cases were generally younger than symptomatic ones, represents a limitation that could bias our comparisons. Given the small size of FTD datasets, balancing our groups by exclusion was not feasible. However, we accounted for age in all our analyses and modeling as a confounding factor and attempted to regress out its impact. By doing so, we aim to minimize potential age-related biases in our results.

Another important limitation is the relatively small number of symptomatic cases, particularly when compared to studies applying DEBM to diseases such as Alzheimer’s. This smaller sample size may contribute to challenges in GMM, including occasional instability in biomarker modeling. Such instability is influenced, in part, by the overlap between normal and abnormal Gaussian distributions, which becomes more pronounced when samples with abnormal biomarker values are limited. However, our study represents one of the largest cohorts available for FTD research, surpassing the size of previous reports and enhancing the reliability and generalizability of our findings.

While many neuroimaging biomarkers could be explored, we focused on a curated list identified in the literature as relevant to FTD. Notably, in DEBM, adding more biomarkers can reduce certainty in the event order, emphasizing the need for deliberate selection.

Our study boasts several strengths that contribute to the robustness of our findings in addition to the larger number of participants compared to previous reports. The quality of our WMH pipeline, which remains robust across multisite data acquisition and functions effectively without the need for FLAIR MRI modality, further solidifies the reliability of our WMH measurements. The application of a robust event-based modeling approach, resilient to missing values, provides a comprehensive understanding of the sequencing of biomarker abnormalities across different mutation groups.

In conclusion, our study not only contributes valuable insights into the distribution and dynamics of WMH in genetic FTD but also highlights the potential role of WMH in refining disease progression models in *GRN* mutations. It will be important to uncover the pathological differences explaining why some *GRN* carriers develop more WMH than others. Further research in this direction may uncover novel avenues for therapeutic interventions targeting neuroinflammatory processes associated with WMH in genetic FTD.

## Data availability

The data used in this study is part of the GENFI dataset and can be accessed upon reasonable request through the study website (www.genfi.org), subject to review and approval by the GENFI data access committee.

## Acknowledgements

We would like to thank all participants and their families for taking part in the GENFI study.

## Abbreviations

FTD: Frontotemporal Dementia
WMHs: White Matter hyperintensities
MRI: Magnetic Resonance Imaging
T1w: T1-weighted
T2w: T2-weighted
NfL: Neurofilament Light chain
GFAP: Glial Fibrillary Acidic Protein
GIF: Geodesic Information Flow
DEBM: Discriminative Event-Based Modeling
GMM: Gaussian Mixture Modeling
bvFTD: behavioral variant of Frontotemporal Dementia
PPA: Primary Progressive Aphasia
ALS: Amyotrophic Lateral Sclerosis
AUC: Area Under the Curve.

## Supplementary material

Supplementary materials are available online.

## Appendix 1

Further details are provided in the Supplementary materials.

## GENFI consortium members

Annabel Nelson, Martina Bocchetta, David Cash, David L Thomas, Emily Todd, Hanya Benotmane, Jennifer Nicholas, Kiran Samra, Rachelle Shafei, Carolyn Timberlake, Thomas Cope, Timothy Rittman, Antonella Alberici, Enrico Premi, Roberto Gasparotti, Valentina Cantoni, Emanuele Buratti, Andrea Arighi, Chiara Fenoglio, Elio Scarpini, Giorgio Fumagalli, Vittoria Borracci, Giacomina Rossi, Giorgio Giaccone, Giuseppe Di Fede, Paola Caroppo, Pietro Tiraboschi, Sara Prioni, Veronica Redaelli, David Tang-Wai, Ekaterina Rogaeva, Miguel Castelo-Branco, Morris Freedman, Ron Keren, Sandra Black, Sara Mitchell, Christen Shoesmith, Robart Bartha, Rosa Rademakers, Janne M. Papma, Lucia Giannini, Rick van Minkelen, Yolande Pijnenburg, Benedetta Nacmias, Camilla Ferrari, Cristina Polito, Gemma Lombardi, Valentina Bessi, Michele Veldsman, Christin Andersson, Hakan Thonberg, Linn Öijerstedt, Vesna Jelic, Paul Thompson, Tobias Langheinrich, Albert Lladó, Anna Antonell, Jaume Olives, Mircea Balasa, Nuria Bargalló, Sergi Borrego-Ecija, Ana Verdelho, Ana Gorostidi, Jorge Villanua, Marta Cañada, Mikel Tainta, Miren Zulaica, Myriam Barandiaran, Patricia Alves, Benjamin Bender, Lisa Graf, Annick Vogels, Mathieu Vandenbulcke, Philip Van Damme, Rose Bruffaerts, Koen Poesen, Pedro Rosa-Neto, Serge Gauthier, Anne Bertrand, Aurélie Funkiewiez, Daisy Rinaldi, Dario Saracino, Olivier Colliot, Sabrina Sayah, Catharina Prix, Elisabeth Wlasich, Olivia Wagemann, Sandra Loosli, Sonja Schönecker, Tobias Hoegen, Jolina Lombardi, Sarah Anderl-Straub, Adeline Rollin, Gregory Kuchcinski, Maxime Bertoux, Thibaud Lebouvier, Vincent Deramecour, Beatriz Santiago, Diana Duro, Maria João Leitão, Maria Rosario Almeida, Miguel Tábuas-Pereira, Sónia Afonso

