## supplementary for "White Matter Hyperintensities Precede other Biomarkers in *GRN* Frontotemporal Dementia"

### Supplementary materials

**Table S1. Average regional WMH volume across distinct mutation cohorts** (raw values in mm3)

| | Non-carriers (Healthy control) | *C9orf72* expansion carriers | | *GRN* mutation carriers | | *MAPT* mutation carriers | |
|---|---|---|---|---|---|---|---|
| | | **Presymptomatic** | **Symptomatic** | **Presymptomatic** | **Symptomatic** | **Presymptomatic** | **Symptomatic** |
| Whole Brain | 6105.097 | 7587.53 | 8672.57 | 6376.52 | 14445.80 | 11181.56 | 16838.93 |
| Left Frontal Lobe | 1556.017 | 1954.209 | 2428.844 | 1562.482 | 4280.289 | 2856.280 | 4219.148 |
| Right Frontal Lobe | 1615.326 | 1928.256 | 2502.688 | 1693.847 | 4369.911 | 3090.220 | 4606.815 |
| Left Temporal Lobe | 423.0705 | 495.488 | 503.558 | 496.686 | 730.667 | 675.960 | 960.259 |
| Right Temporal Lobe | 457.611 | 505.318 | 569.013 | 505.431 | 794.089 | 739.140 | 1166.222 |
| Left Parietal Lobe | 776.164 | 1083.628 | 1038.623 | 790.226 | 1579.578 | 1549.020 | 2530.407 |
| Right Parietal Lobe | 720.255 | 989.853 | 903.156 | 742.693 | 1609.022 | 1359.740 | 2224.037 |
| Left Occipital Lobe | 222.762 | 250.512 | 369.779 | 216.474 | 513.267 | 340.480 | 571.926 |
| Right Occipital Lobe | 229.899 | 260.659 | 315.935 | 238.401 | 430.689 | 364.200 | 387.778 |

**Table S2. Average regional WMH volume across distinct mutation cohorts, with adjustments for age, sex, and scanner site.** The values have been subjected to a log transformation.

| | Non-carriers (Healthy control) | *C9orf72* expansion carriers | | *GRN* mutation carriers | | *MAPT* mutation carriers | |
|---|---|---|---|---|---|---|---|
| | | Presymptomatic | Symptomatic | Presymptomatic | Symptomatic | Presymptomatic | Symptomatic |
| Whole Brain | 0.0000 | 0.0705 | 0.0418 | 0.0195 | 0.2368 | 0.1135 | 0.0365 |
| Left Frontal Lobe | 0.0000 | 0.0768 | 0.0664 | 0.0348 | 0.3110 | 0.1292 | 0.0497 |
| Right Frontal Lobe | 0.0000 | 0.0608 | 0.0687 | 0.0299 | 0.2448 | 0.1329 | 0.0529 |
| Left Temporal Lobe | -0.0000 | 0.0730 | -0.0252 | 0.0111 | 0.0999 | 0.0681 | 0.0543 |
| Right Temporal Lobe | 0.0000 | 0.0301 | -0.0028 | 0.0179 | 0.8669 | 0.0563 | 0.0470 |
| Left Parietal Lobe | 0.0000 | 0.0834 | 0.0160 | -0.0319 | 0.1203 | 0.1113 | -0.0099 |
| Right Parietal Lobe | 0.0000 | 0.11076 | -0.0001 | 0.0264 | 0.1831 | 0.1792 | 0.0955 |
| Left Occipital Lobe | 0.0000 | 0.1300 | -0.0991 | -0.0211 | 0.1414 | 0.1761 | 0.0959 |
| Right Occipital Lobe | -0.0000 | 0.0682 | -0.0722 | 0.0040 | -0.0957 | 0.1651 | -0.0274 |

**Table S3. Comparison of adjusted WMHs across brain lobes among different mutation groups.** Only comparisons with unadjusted p-values ≤ 0.05 are shown. WMHs are adjusted for age, sex, and scanner site.

| Region | Mutation types | Unadjusted p-value | Bonferroni Adjusted p-value | FDR Adjusted p-value |
|---|---|---|---|---|
| Whole Brain | Symptomatic *GRN* > control | 0.0001 | 0.0006 | 0.0006 |
| Left Frontal lobe | Symptomatic *GRN* > Symptomatic *C9orf72* | 0.0017 | 0.0103 | 0.0051 |
| Left Frontal lobe | Symptomatic *GRN* > control | < 0.0001 | < 0.0001 | < 0.0001 |
| Left Frontal lobe | Symptomatic *GRN* > Symptomatic *MAPT* | 0.0042 | 0.0253 | 0.0084 |
| Right Frontal lobe | Symptomatic *GRN* > control | 0.0005 | 0.0030 | 0.0030 |
| Whole Brain | Symptomatic *GRN* > Symptomatic *C9orf72* | 0.0165 | 0.0992 | 0.0496 |
| Whole Brain | Symptomatic *GRN* > Symptomatic *MAPT* | 0.0246 | 0.1474 | 0.0491 |
| Right Frontal lobe | Symptomatic *GRN* > Symptomatic *MAPT* | 0.0522 | 0.3131 | 0.1565 |
| Left Frontal lobe | Presymptomatic *C9orf72* > control | 0.0488 | 0.2929 | 0.2929 |
| Left Temporal | Presymptomatic *C9orf72* > control | 0.0320 | 0.1918 | 0.1918 |
| Left Temporal | Symptomatic *GRN* > control | 0.0530 | 0.3180 | 0.3180 |
| Left Occipital | Presymptomatic *C9orf72* > control | 0.0295 | 0.1770 | 0.1770 |
| Left Occipital | Presymptomatic *MAPT* > control | 0.0481 | 0.2888 | 0.1444 |

**Table S4. Comparison of adjusted WMHs across brain lobes among different mutation groups in different disease stages.**

| Region | Genetic Group | Genetic status | Unadjusted p-value | Bonferroni Adjusted p-value | FDR Adjusted p-value |
|---|---|---|---|---|---|
| Left Frontal | all | Symptomatic > control | 0.0002 | 0.0005 | 0.0005 |
| Right Frontal | all | Symptomatic > control | 0.0025 | 0.0075 | 0.0075 |
| Whole brain | all | Symptomatic > control | 0.0039 | 0.0116 | 0.0116 |
| Whole brain | *GRN* | Symptomatic > Presymptomatic | 0.0005 | 0.00156 | 0.0008 |
| Right Frontal | *GRN* | Symptomatic > Presymptomatic | 0.0031 | 0.0094 | 0.0047 |
| Left Frontal | *GRN* | Symptomatic > Presymptomatic | < 0.0001 | < 0.0001 | < 0.0001 |
| Left Frontal | all | Symptomatic > presymptomatic | 0.0281 | 0.0843 | 0.0422 |
| Left Frontal | all | Presymptomatic > control | 0.0485 | 0.1455 | 0.0485 |
| Right Occipital | *C9orf72* | Presymptomatic > Symptomatic | 0.0416 | 0.1248 | 0.0624 |
| Right occipital | all | Presymptomatic > Symptomatic | 0.0322 | 0.0968 | 0.0968 |
| Left Occipital | all | Presymptomatic > control | 0.0571 | 0.1713 | 0.1713 |

**Table S5. Mean Square Error (MSE) of Gaussian Mixture Modeling in the DEBM Framework.** This table shows the MSE values for distributions fitted using Gaussian mixture modeling as part of the DEBM model. The MSE is calculated by comparing the fitted Gaussian mixture model with the histogram of the data. Biomarkers with an MSE exceeding 15 are considered poorly fitted, excluded from DEBM analysis, and included in the longitudinal analysis instead.

| Biomarker | Mean Square Error |
|---|---|
| NfL | 9.8408 |
| GFAP | 4.5165 |
| Total WMH | 11.5816 |
| WMH Frontal | 10.2852 |
| WMH Temporal | 6.8822 |
| Ventricle | 12.9720 |
| Frontal GM | 9.5862 |
| Temporal GM | 6.7830 |
| Cerebellum | 7.2356 |
| Insula | 67.4482 |
| Basal Ganglia | 91.9080 |
| Thalamus | 85.7825 |
| Amygdala | 174.4104 |
| Hippocampus | 77.6946 |
| Cingulate | 66.2930 |

**Table S6. Comparison of raw biomarker values between FTD and healthy controls,** without adjustment for age and sex, to identify statistically significant differences as part of the inclusion criteria for DEBM analysis.

| Biomarker | tStat | pVal |
|---|---|---|
| NfL | -15.025 | < 0.0001 |
| GFAP | -7.664 | < 0.0001 |
| Total WMH | -6.338 | < 0.0001 |
| WMH Frontal | -6.987 | < 0.0001 |
| WMH Parietal | -4.918 | < 0.0001 |
| Ventricle | -14.723 | < 0.0001 |
| Frontal GM | -11.606 | < 0.0001 |
| Temporal GM | -8.853 | < 0.0001 |
| Cerebellum | -5.581 | < 0.0001 |
| Insula | -15.169 | < 0.0001 |
| Basal Ganglia | -8.337 | < 0.0001 |
| Thalamus | -6.695 | < 0.0001 |
| Amygdala | -4.858 | < 0.0001 |
| Hippocampus | -6.214 | < 0.0001 |
| Cingulate | -11.128 | < 0.0001 |

**Table S7. Correlation between DEBM staging and clinical scores (among mutation carriers),** including Mini-Mental State Examination (MMSE), Trail Making Test Part B (TMTB) time, Digit Symbol substitution test, Boston naming, and Verbal Fluency (VF) combined score.

| Clinical score | Spearman's Rank Correlation | t | df | p-value |
|---|---|---|---|---|
| MMSE | -0.48 | -6.99 | 164 | < 0.001 |
| TMTB time | 0.36 | 4.81 | 160 | < 0.001 |
| Digit symbol | -0.43 | -6.22 | 168 | < 0.001 |
| Boston naming | -0.28 | -3.88 | 171 | < 0.001 |
| VF combined | -0.40 | -5.57 | 165 | < 0.001 |

**Table S8. Parameter estimates for the longitudinal analysis model,** where the change in subcortical regions over time (ΔResponse biomarker/Δt) is modeled as a function of baseline WMH (predictor biomarker), age, sex, neurofilament light chain (NfL), education, and baseline subcortical biomarker.

| **Predictor biomarker** | **Response biomarker** | **Predictor biomarker baseline** | | | **Response biomarker baseline** | | **NfL** | | **Age** | | **Sex** | | **Education** | |
|---|---|---|---|---|---|---|---|---|---|---|---|---|---|---|
| | | **tStat** | **pVal** | **FDR pVal** | **tStat** | **pVal** | **tStat** | **pVal** | **tStat** | **pVal** | **tStat** | **pVal** | **tStat** | **pVal** |
| Total WMH | Thalamus | -0.68 | 0.497 | 0.540193 | -2.76 | 0.007 | -2.59 | 0.011 | -0.39 | 0.699 | -1.58 | 0.119 | -0.44 | 0.661 |
| Total WMH | Basal Ganglia | -1.81 | 0.074 | 0.111 | -1.19 | 0.239 | -4.49 | < 0.001 | 1.54 | 0.127 | -0.11 | 0.915 | -1.19 | 0.236 |
| Total WMH | Amygdala | -2.80 | 0.006 | **0.038** | -3.81 | < 0.001 | -2.11 | 0.038 | -1.07 | 0.288 | 0.44 | 0.660 | -0.42 | 0.675 |
| Total WMH | Hippocampus | -2.32 | 0.023 | **0.050** | -3.80 | < 0.001 | -2.60 | 0.011 | -0.88 | 0.382 | 0.87 | 0.387 | -1.27 | 0.207 |
| Total WMH | Cingulate | -2.28 | 0.025 | **0.050** | -1.71 | 0.090 | -3.29 | 0.001 | 1.12 | 0.264 | -0.51 | 0.607 | -0.54 | 0.588 |
| Total WMH | Insula | 0.61 | 0.540 | 0.540 | 0.003 | 0.997 | -5.60 | < 0.001 | 2.30 | 0.024 | 1.61 | 0.111 | -1.33 | 0.186 |

**Table S9. Parameter estimates for the longitudinal analysis model,** where the change in WMH over time (ΔResponse biomarker/Δt) is modeled as a function of baseline predictor biomarker (subcortical volumes), age, sex, neurofilament light chain (NfL), education, and baseline WMH.

| **Predictor biomarker** | **Response biomarker** | **Predictor biomarker baseline** | | | **Response biomarker baseline** | | **NfL** | | **Age** | | **Sex** | | **Education** | |
|---|---|---|---|---|---|---|---|---|---|---|---|---|---|---|
| | | **tStat** | **pVal** | **FDR pVal** | **tStat** | **pVal** | **tStat** | **pVal** | **tStat** | **pVal** | **tStat** | **pVal** | **tStat** | **pVal** |
| Thalamus | Total WMH | 1.67 | 0.098 | 0.497 | -1.47 | 0.144 | 1.93 | 0.057 | 0.88 | 0.380 | 0.95 | 0.345 | 0.92 | 0.359 |
| Basal Ganglia | Total WMH | -0.03 | 0.974 | 0.991 | -1.99 | 0.049 | 1.57 | 0.120 | 0.42 | 0.679 | 0.51 | 0.613 | 1.16 | 0.251 |
| Amygdala | Total WMH | 1.40 | 0.166 | 0.497 | -1.94 | 0.056 | 1.76 | 0.083 | 0.86 | 0.394 | 0.50 | 0.615 | 0.88 | 0.378 |
| Hippocampus | Total WMH | 0.01 | 0.991 | 0.991 | -1.96 | 0.054 | 1.68 | 0.097 | 0.41 | 0.680 | 0.52 | 0.606 | 1.14 | 0.258 |
| Cingulate | Total WMH | 0.28 | 0.781 | 0.991 | -1.91 | 0.060 | 1.72 | 0.088 | 0.46 | 0.648 | 0.56 | 0.577 | 1.15 | 0.255 |
| Insula | Total WMH | -0.34 | 0.732 | 0.991 | -2.06 | 0.043 | 1.44 | 0.154 | 0.34 | 0.735 | 0.43 | 0.668 | 1.18 | 0.243 |

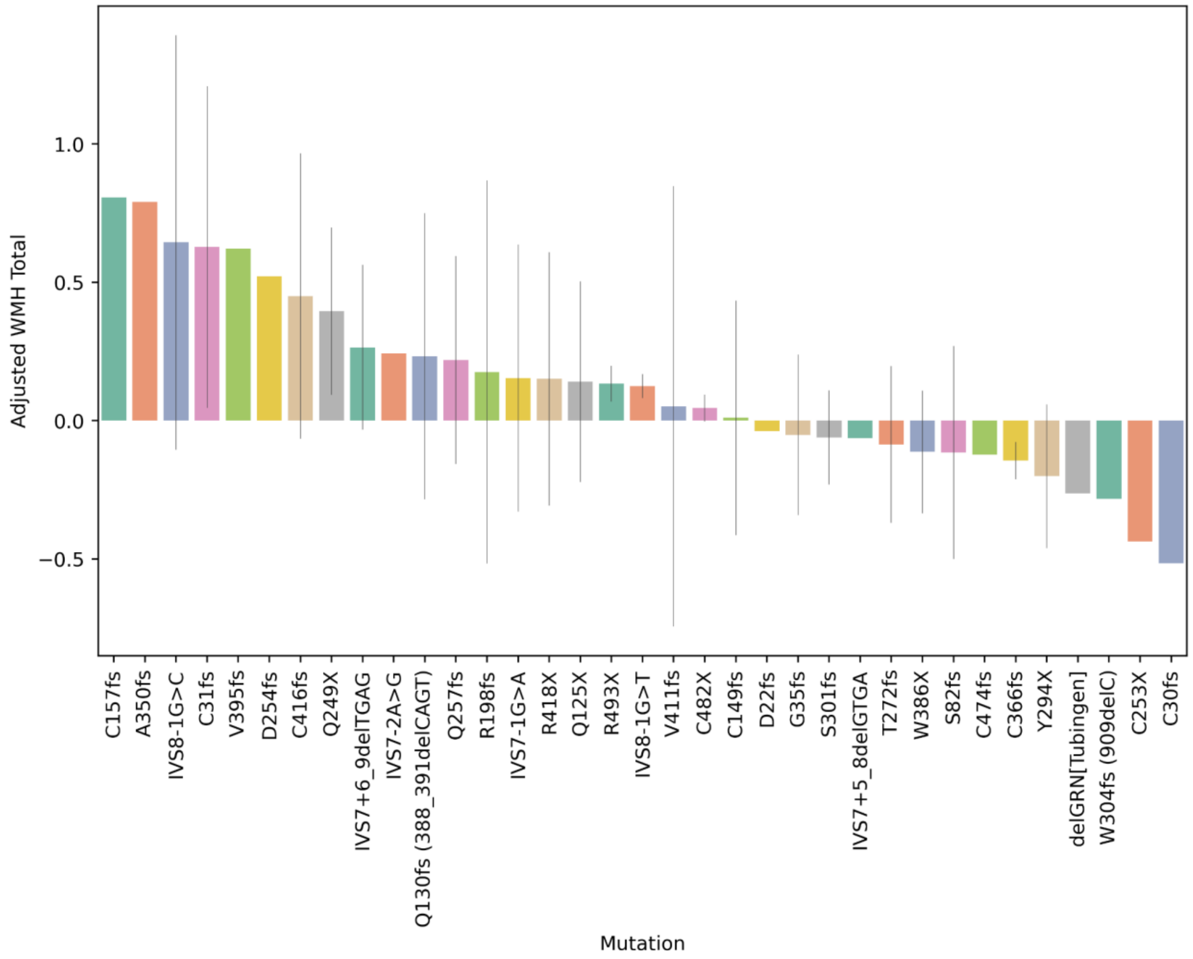


**Figure S1. Adjusted WMH Volumes by GRN Mutation Subtype** Each bar represents the average of adjusted WMH volumes (controlled for age, sex, and scanner site as explained in Equation 1) in different GRN mutation subtypes among all GRN carriers (both presymptomatic and symptomatic).

#### GENFI consortium members

Annabel Nelson Department of Neurodegenerative Disease, Dementia Research Centre, UCL Queen Square Institute of Neurology, London, UK; Martina Bocchetta Department of Neurodegenerative Disease, Dementia Research Centre, UCL Queen Square Institute of Neurology, London, UK; David Cash Department of Neurodegenerative Disease, Dementia Research Centre, UCL Queen Square Institute of Neurology, London, UK; David L Thomas Neuroimaging Analysis Centre, Department of Brain Repair and Rehabilitation, UCL Institute of Neurology, Queen Square, London, UK; Emily Todd Department of Neurodegenerative Disease, Dementia Research Centre, UCL Queen Square Institute of Neurology, London, UK;

Hanya Benotmane UK Dementia Research Institute at University College London, UCL Queen Square Institute of Neurology, London, UK; Jennifer Nicholas Department of Medical Statistics, London School of Hygiene and Tropical Medicine, London, UK; Kiran Samra Department of Neurodegenerative Disease, Dementia Research Centre, UCL Queen Square Institute of Neurology, London, UK; Rachelle Shafei Department of Neurodegenerative Disease, Dementia Research Centre, UCL Queen Square Institute of Neurology, London, UK; Carolyn Timberlake Department of Clinical Neurosciences, University of Cambridge, Cambridge, UK; Thomas Cope Department of Clinical Neuroscience, University of Cambridge, Cambridge, UK; Timothy Rittman Department of Clinical Neurosciences, University of Cambridge, Cambridge, UK; Antonella Alberici Centre for Neurodegenerative Disorders, University of Brescia, Brescia, Italy; Enrico Premi Stroke Unit, ASST Brescia Hospital, Brescia, Italy; Roberto Gasparotti Neuroradiology Unit, University of Brescia, Brescia, Italy; Valentina Cantoni Centre for Neurodegenerative Disorders, Department of Clinical and Experimental Sciences, University of Brescia, Brescia, Italy; Emanuele Buratti ICGEB, Trieste, Italy; Andrea Arighi Fondazione IRCCS Ca' Granda Ospedale Maggiore Policlinico, Neurodegenerative Diseases Unit, Milan, Italy; University of Milan, Centro Dino Ferrari, Milan, Italy; Chiara Fenoglio Fondazione IRCCS Ca' Granda Ospedale Maggiore Policlinico, Neurodegenerative Diseases Unit, Milan, Italy; University of Milan, Centro Dino Ferrari, Milan, Italy; Elio Scarpini Fondazione IRCCS Ca' Granda Ospedale Maggiore Policlinico, Neurodegenerative Diseases Unit, Milan, Italy; University of Milan, Centro Dino Ferrari, Milan, Italy; Giorgio Fumagalli Fondazione IRCCS Ca' Granda Ospedale Maggiore Policlinico, Neurodegenerative Diseases Unit, Milan, Italy; University of Milan, Centro Dino Ferrari, Milan, Italy; Vittoria Borracci Fondazione IRCCS Ca' Granda Ospedale Maggiore Policlinico, Neurodegenerative Diseases Unit, Milan, Italy; University of Milan, Centro Dino Ferrari, Milan, Italy; Giacomina Rossi Fondazione IRCCS Istituto Neurologico Carlo Besta, Milano, Italy; Giorgio Giaccone Fondazione IRCCS Istituto Neurologico Carlo Besta, Milano, Italy; Giuseppe Di Fede Fondazione IRCCS Istituto Neurologico Carlo Besta, Milano, Italy; Paola Caroppo Fondazione IRCCS Istituto Neurologico Carlo Besta, Milano, Italy; Pietro Tiraboschi Fondazione IRCCS Istituto Neurologico Carlo Besta, Milano, Italy; Sara Prioni Fondazione IRCCS Istituto Neurologico Carlo Besta, Milano, Italy; Veronica Redaelli Fondazione IRCCS Istituto Neurologico Carlo Besta, Milano, Italy; David Tang-Wai The University Health Network, Krembil Research Institute, Toronto, Canada; Ekaterina Rogaeva Tanz Centre for Research in Neurodegenerative Diseases, University of Toronto, Toronto, Canada; Miguel Castelo-Branco Faculty of Medicine, University of Coimbra, Coimbra,

Portugal; Morris Freedman Baycrest Health Sciences, Rotman Research Institute, University of Toronto, Toronto, Canada; Ron Keren The University Health Network, Toronto Rehabilitation Institute, Toronto, Canada; Sandra Black Sunnybrook Health Sciences Centre, Sunnybrook Research Institute, University of Toronto, Toronto, Canada; Sara Mitchell Sunnybrook Health Sciences Centre, Sunnybrook Research Institute, University of Toronto, Toronto, Canada; Christen Shoesmith Department of Clinical Neurological Sciences, University of Western Ontario, London, Ontario, Canada; Robart Bartha Department of Medical Biophysics, The University of Western Ontario, London, Ontario, Canada; Centre for Functional and Metabolic Mapping, Robarts Research Institute, The University of Western Ontario, London, Ontario, Canada; Rosa Rademakers Center for Molecular Neurology, University of Antwerp Jackie Poos Department of Neurology, Erasmus Medical Center, Rotterdam, Netherlands; Janne M. Papma Department of Neurology, Erasmus Medical Center, Rotterdam, Netherlands; Lucia Giannini Department of Neurology, Erasmus Medical Center, Rotterdam, Netherlands; Rick van Minkelen Department of Clinical Genetics, Erasmus Medical Center, Rotterdam, Netherlands; Yolande Pijnenburg Amsterdam University Medical Centre, Amsterdam VUmc, Amsterdam, Netherlands; Camilla Ferrari Department of Neuroscience, Psychology, Drug Research and Child Health, University of Florence, Florence, Italy; Enrico Fainardi Neuroradiology Unit, Department of Experimental and Clinical Biomedical Sciences, University of Florence, Florence, Italy; Stefano Chiti Neuroradiology Unit, Department of Experimental and Clinical Biomedical Sciences, University of Florence, Florence, Italy; Giulia Giacomucci Department of Neuroscience, Psychology, Drug Research and Child Health, University of Florence, Florence, Italy; Valentina Moschini Neurology unit , Careggi university Hospital, Florence Italy; Valentina Bessi Department of Neuroscience, Psychology, Drug Research and Child Health, University of Florence, Florence, Italy; Michele Veldsman Nuffield Department of Clinical Neurosciences, Medical Sciences Division, University of Oxford, Oxford, UK; Christin Andersson Department of Clinical Neuroscience, Karolinska Institutet, Stockholm, Sweden; Hakan Thonberg Center for Alzheimer Research, Division of Neurogeriatrics, Karolinska Institutet, Stockholm, Sweden; Linn Öijerstedt Center for Alzheimer Research, Division of Neurogeriatrics, Department of Neurobiology, Care Sciences and Society, Bioclinicum, Karolinska Institutet, Solna, Sweden; Unit for Hereditary Dementias, Theme Aging, Karolinska University Hospital, Solna, Sweden; Vesna Jelic Division of Clinical Geriatrics, Karolinska Institutet, Stockholm, Sweden; Paul Thompson Division of Neuroscience and Experimental Psychology, Wolfson Molecular Imaging Centre, University of Manchester, Manchester, UK; Tobias Langheinrich Division of Neuroscience and Experimental Psychology, Wolfson Molecular Imaging Centre, University of Manchester, Manchester, UK; Manchester Centre for Clinical Neurosciences, Department of Neurology, Salford Royal NHS Foundation Trust, Manchester, UK; Albert Lladó Alzheimer's disease and Other Cognitive Disorders Unit, Neurology Service, Hospital Clínic, Barcelona, Spain; Anna Antonell Alzheimer's disease and Other Cognitive Disorders Unit, Neurology Service, Hospital Clínic, Barcelona, Spain; Jaume Olives Alzheimer's disease and Other Cognitive Disorders Unit, Neurology Service, Hospital Clínic, Barcelona, Spain; Mircea Balasa Alzheimer's disease and Other Cognitive Disorders Unit, Neurology Service, Hospital Clínic, Barcelona, Spain; Nuria Bargalló Imaging Diagnostic Center, Hospital Clínic, Barcelona, Spain; Sergi Borrego-Ecija Alzheimer's disease and Other Cognitive Disorders Unit, Neurology Service, Hospital Clínic, Barcelona, Spain; Ana Verdelho Department of Neurosciences and Mental Health, Centro Hospitalar Lisboa Norte - Hospital de Santa Maria Neuroscience Area, Biodonostia Health Research Insitute, San Sebastian, Gipuzkoa, Spain; Ana Gorostidi Neuroscience Area, Biodonostia Health Research Insitute, San Sebastian, Gipuzkoa, Spain; Jorge Villanua OSATEK, University of Donostia, San Sebastian, Gipuzkoa, Spain; Marta Cañada CITA Alzheimer, San Sebastian, Gipuzkoa, Spain; Mikel Tainta Neuroscience Area, Biodonostia Health Research Insitute, San Sebastian, Gipuzkoa, Spain; Miren Zulaica Neuroscience Area, Biodonostia Health Research Insitute, San Sebastian, Gipuzkoa, Spain; Myriam Barandiaran Cognitive Disorders Unit, Department of Neurology, Donostia University Hospital, San Sebastian, Gipuzkoa, Spain; Neuroscience Area, Biodonostia Health Research Insitute, San Sebastian, Gipuzkoa, Spain; Patricia Alves Neuroscience Area, Biodonostia Health Research Insitute, San Sebastian, Gipuzkoa, Spain; Department of Educational Psychology and Psychobiology, Faculty of Education, International University of La Rioja, Logroño, Spain; Benjamin Bender Department of Diagnostic and Interventional Neuroradiology, University of Tübingen, Tübingen, Germany; Lisa Graf Department of Neurodegenerative Diseases, Hertie-Institute for Clinical Brain Research and Center of Neurology, University of Tübingen, Tübingen, Germany; Annick Vogels Department of Human Genetics, KU Leuven, Leuven, Belgium; Mathieu Vandenbulcke Geriatric Psychiatry Service, University Hospitals Leuven, Belgium; Neuropsychiatry, Department of Neurosciences, KU Leuven, Leuven, Belgium; Philip Van Damme Neurology Service, University Hospitals Leuven, Belgium; Laboratory for Neurobiology, VIB-KU Leuven Centre for Brain Research, Leuven, Belgium; Rose Bruffaerts Department of Biomedical Sciences, University of Antwerp, Antwerp, Belgium; Biomedical Research Institute, Hasselt University, 3500 Hasselt, Belgium; Koen Poesen Laboratory for Molecular

Neurobiomarker Research, KU Leuven, Leuven, Belgium; Pedro Rosa-Neto Translational Neuroimaging Laboratory, McGill Centre for Studies in Aging, McGill University, Montreal, Québec, Canada; Serge Gauthier Alzheimer Disease Research Unit, McGill Centre for Studies in Aging, Department of Neurology Reference Network for Rare Neurological Diseases (ERN-RND)Anne Bertrand Sorbonne Université, Paris Brain Institute – Institut du Cerveau – ICM, Inserm U1127, CNRS UMR 7225, AP-HP - Hôpital Pitié-Salpêtrière, Paris, France; Inria, Aramis project-team, F-75013, Paris, France; Centre pour l'Acquisition et le Traitement des Images, Institut du Cerveau et la Moelle, Paris, France; Aurélie Funkiewiez Centre de référence des démences rares ou précoces, IM2A, Département de Neurologie, AP-HP - Hôpital Pitié-Salpêtrière, Paris, France; Sorbonne Université, Paris Brain Institute – Institut du Cerveau – ICM, Inserm U1127, CNRS UMR 7225, AP-HP - Hôpital Pitié-Salpêtrière, Paris, France; Daisy Rinaldi Centre de référence des démences rares ou précoces, IM2A, Département de Neurologie, AP-HP - Hôpital Pitié-Salpêtrière, Paris, France; Sorbonne Université, Paris Brain Institute – Institut du Cerveau – ICM, Inserm U1127, CNRS UMR 7225, AP-HP - Hôpital Pitié-Salpêtrière, Paris, France; Département de Neurologie, AP-HP - Hôpital Pitié-Salpêtrière, Paris, France; Dario Saracino Sorbonne Université, Paris Brain Institute – Institut du Cerveau – ICM, Inserm U1127, CNRS UMR 7225, AP-HP - Hôpital Pitié-Salpêtrière, Paris, France; Inria, Aramis project-team, F-75013, Paris, France; Centre de référence des démences rares ou précoces, IM2A, Département de Neurologie, AP-HP - Hôpital Pitié-Salpêtrière, Paris, France; Olivier Colliot Sorbonne Université, Paris Brain Institute – Institut du Cerveau – ICM, Inserm U1127, CNRS UMR 7225, AP-HP - Hôpital Pitié-Salpêtrière, Paris, France; Inria, Aramis project team, F-75013, Paris, France; Centre pour l'Acquisition et le Traitement des Images, Institut du Cerveau et la Moelle, Paris, France; Sabrina Sayah Sorbonne Université, Paris Brain Institute – Institut du Cerveau – ICM, Inserm U1127, CNRS UMR 7225, AP-HP - Hôpital Pitié-Salpêtrière, Paris, France; Catharina Prix Neurologische Klinik, Ludwig-Maximilians-Universität München, Munich, Germany; Elisabeth Wlasich Neurologische Klinik, Ludwig-Maximilians-Universität München, Munich, Germany; Olivia Wagemann Neurologische Klinik, Ludwig-Maximilians-Universität München, Munich, Germany; Sandra Loosli Neurologische Klinik, Ludwig-Maximilians Universität München, Munich, Germany; Sonja Schönecker Neurologische Klinik, Ludwig-Maximilians-Universität München, Munich, Germany; Tobias Hoegen Neurologische Klinik, Ludwig-Maximilians-Universität München, Munich, Germany; Jolina Lombardi Department of Neurology, University of Ulm, Ulm; Sarah Anderl-Straub Department of Neurology, University of Ulm, Ulm, Germany; Adeline Rollin CHU, CNR-MAJ, Labex Distalz, LiCEND Lille, France; Gregory Kuchcinski Univ Lille,

France; Inserm 1172, Lille, France; CHU, CNR-MAJ, Labex Distalz, LiCEND Lille, France; Maxime Bertoux Inserm 1172, Lille, France; CHU, CNR-MAJ, Labex Distalz, LiCEND Lille, France; Thibaud Lebouvier Univ Lille, France; Inserm 1172, Lille, France; CHU, CNR-MAJ, Labex Distalz, LiCEND Lille, France; Vincent Deramecourt Univ Lille, France; Inserm 1172, Lille, France; CHU, CNR-MAJ, Labex Distalz, LiCEND Lille, France; Beatriz Santiago Neurology Department, Centro Hospitalar e Universitario de Coimbra, Coimbra, Portugal; Diana Duro Faculty of Medicine, University of Coimbra, Coimbra, Portugal; Maria João Leitão Centre of Neurosciences and Cell Biology, Universidade de Coimbra, Coimbra, Portugal; Maria Rosario Almeida Faculty of Medicine, University of Coimbra, Coimbra, Portugal; Miguel Tábuas-Pereira Neurology Department, Centro Hospitalar e Universitario de Coimbra, Coimbra, Portugal; Sónia Afonso Instituto Ciencias Nucleares Aplicadas a Saude, Universidade de Coimbra, Coimbra, Portugal
